# Highly pathogenic avian influenza H5N1 virus infections in pinnipeds and seabirds in Uruguay: a paradigm shift to virus transmission in South America

**DOI:** 10.1101/2023.12.14.571746

**Authors:** Gonzalo Tomás, Ana Marandino, Yanina Panzera, Sirley Rodríguez, Gabriel Luz Wallau, Filipe Zimmer Dezordi, Ramiro Pérez, Lucía Bassetti, Raúl Negro, Joaquín Williman, Valeria Uriarte, Fabiana Grazioli, Carmen Leizagoyen, Sabrina Riverón, Jaime Coronel, Soledad Bello, Enrique Páez, Martín Lima, Virginia Méndez, Ruben Pérez

## Abstract

The highly pathogenic avian influenza viruses of the clade 2.3.4.4b have caused unprecedented deaths in South American wild birds, poultry, and marine mammals. In September 2023, pinnipeds and seabirds appeared dead on the Uruguayan Atlantic coast. Sixteen influenza virus strains were characterized by real-time reverse transcription PCR and genome sequencing in samples from sea lions (*Otaria flavescens*), fur seals (*Arctocephalus australis*), and terns (*Sterna hirundinacea*). Phylogenetic and ancestral reconstruction analysis showed that these strains have pinnipeds as the most likely ancestral host, representing a recent introduction of the clade 2.3.4.4b in Uruguay. The Uruguayan and closely related strains from Peru (sea lions) and Chile (sea lions and a human case) carry mammalian adaptative residues 591K and 701N in the viral polymerase basic protein 2 (PB2). Our findings suggest that the clade 2.3.4.4b strains in South America may have spread from mammals to mammals and seabirds, revealing a new transmission route.

## Introduction

*The Alphainfluenzavirus influenzae* species (family *Orthomyxoviridae*) is a prominent pathogen in birds and mammals. The virus is divided into subtypes based on the genetic and antigenic properties of the hemagglutinin (HA) and neuraminidase (NA) (1).

Migratory wild birds, especially waterfowl, are the natural host and reservoir of avian influenza virus (AIV) and harbor most combinations of circulating subtypes (2). Influenza in mammals evolved from the adaptation of ancestral avian viruses. The remarkable ability of the influenza virus to adapt to new hosts is associated with its segmented negative single-stranded RNA genome, which mutates and reassorts easily (1).

Low pathogenicity AIV (LPAIV) of the H5 and H7 subtypes may evolve towards highly pathogenic AIV (HPAIV) upon transmission into highly dense domestic bird populations (3). HPAIV strains are primarily characterized by multiple basic amino acid residues at the HA cleavage site processed by ubiquitous proteases, resulting in a fast-spreading deadly disease with increased pathogenicity (4).

Since October 2020, the HPAIV clade 2.3.4.4b (H5N1) from the Goose/Guangdong (Gs/GD) lineage has been responsible for an unprecedented number of deaths in wild birds and poultry (5). The clade 2.3.4.4b spread from Asia to Africa and Europe and reached North America via Iceland (6). The first detections of HPAIV clade 2.3.4.4b in North America were in wild and domestic birds in November 2021 in Canada and late December 2021 in the United States (7–9). In October 2022, it emerged in Latin America, with severe outbreaks identified in Chile, Colombia, Ecuador, Mexico, Peru, and Venezuela (10–12). In Uruguay, HPAIV H5N1 was detected in wild birds and backyard poultry from February to May 2023 (13).

There have been increasing reports of deadly outbreaks among wild and captive mammals caused by clade 2.3.4.4b (12,14–18). Both terrestrial and marine mammals have been affected, including farmed minks (*Neovison vison*) in Spain, true seals (*Phocidae*) in the United States of America, and sea lions (*Otaria flavescens/byronia*) in Peru and Chile (12,14,16,19,20). Several countries have reported infected domestic animals, including cats and dogs (21).

The high frequency of HPAIV spillover into mammals, including humans, raises serious concerns about a potential global pandemic as the virus could adapt and transmit among mammals. However, reporting HPAIV H5N1 in mammals was primarily associated with bird-to-mammal transmission (21).

Marine mammals seem particularly susceptible to avian influenza. In Peru and Chile, more than 15,000 infected sea lions died (12,19,20). The southernmost case of H5N1 on the Pacific coast was detected in June 2023 in a sea lion from Puerto Williams, Chile. The first infected sea lions on the Atlantic coast were reported in the southernmost tip of Argentina (Rio Grande, Tierra del Fuego Province) in August 2023 (22).

The present study conducted viral detection and genome sequencing on infected sea lions (*Otaria flavescens*), fur seals (*Arctocephalus australis*), and sea birds (*Sterna hirundinacea)* sampled in Uruguay from September to October 2023. Our findings suggest mammal-to-mammal transmission and bird infection by viruses of mammalian origin.

## MATERIALS AND METHODS

### AIV Detection and HA Subtyping

During September and early October 2023, ocular, oropharyngeal, encephalic, rectal/cloacal, and fecal swabs were obtained from pinnipeds and seabirds from different regions of the Uruguayan coastline. Samples were collected and promptly transported for molecular diagnosis (13). Clinical signs could not be determined in deceased animals. Carcasses were buried immediately according to sanitary protocol, precluding autopsy.

Viral detection and HA subtyping were performed by real-time reverse transcription PCR, as previously described (13).

### Illumina Sequencing

Complementary DNA (cDNA) was synthesized using the Superscript II First-Strand Synthesis System (Thermo Fisher Scientific, Waltham, MA, USA). Full AIV segment amplification was performed in a single reaction with a primer pair (23) or using unique primers for each viral segment (24). Amplicons were pooled and purified using AMPure XP (Beckman Coulter, Indianapolis, IN, USA), and 100 ng were subjected to the Nextera™ DNA Flex Library Preparation kit (Illumina, San Diego, CA, USA) employing a unique dual indexing (XT-Index Kit V2-Set A). Libraries quantification and sequencing were performed as previously reported (13).

### Genome analysis

The raw sequencing data was trimmed and filtered using BBDuk and Minimap2 plugins in Geneious (25). The clean reads were mapped to an avian influenza genome.

Assemblies were visually inspected and optimized; annotations were transferred from reference strains and manually curated. Recombinant segments were evaluated with the complete genome datasets using the RDP4 v4.95 software (26).

Intra-host variants were analyzed using Geneious by assembling clean reads to obtain consensus and recording minor single-nucleotide variants (SNV) with a frequency greater than 10% and a minimum coverage depth of 100.

### Dataset

We downloaded all sequences from Ecuador, Peru, Chile, Argentina, and Uruguay from GISAID (https://gisaid.org/) (Oct 7, 2023). The USA strain (A/fox/Minnesota/22/016487/001|EPI_ISL_15078249) was included as an outgroup. Sequences were aligned using MAFFT v 7 (27) and trimmed to each segment’s starting ATG and ending STOP codon. Maximum-likelihood phylogenetic trees were obtained using FastTree (28) and IQTREE 2.2.0 (29) with approximate likelihood ratio tests (aLRT) for internal node support.

### Molecular clock and ancestral state reconstruction

Concatenated segments were used to reconstruct the phylogenetic trees. The root-to-tip correlation found in the best ML tree recovered by IQTREE was R = 0,909 based on TempEst v 1.5.3 (30). The timetree reconstruction was performed using the uncorrelated molecular clock, and three coalescent models were tested using the path sampling analysis (constant size, exponential growth, and Bayesian skygrid). The nucleotide substitution model was set as GTR + G + I, based on the best model found by ModelFinder from IQTREE 2.2.0 in Beast 1.10.5 (31). Base frequencies were estimated from the alignment, and 4 gamma categories were added as parameters. The uncorrelated molecular clock model and Bayesian skygrid coalescent model output the best likelihood results, which were selected for further analysis. ESS values were all above 200 for the tested runs.

Bayesian phylogenetic reconstruction was performed using the Beast 1.10.5 software in the same dataset using the stamped tip dates information and three categorical information for discrete phylogeographic analysis: country, host, and the presence of amino acid substitutions. We set up 500 million chains and sampled every 50,000 trees for enough parameter mixing and estimates. ESS values were above 200, validating the optimal parameter space exploration. Burning was set to 10% of all trees generated.

Finally, a phylogenetic tree was depicted with Figtree. We used Phylopic for silhouette imagens obtention (https://www.phylopic.org/).

### Map and case dynamics through time visualization

To display the distribution of H5N1 HPAI cases in South America associated or not with genomic sequences, we downloaded the dataset from https://empres-i.apps.fao.org/. We set up Microreact instances harboring the whole dataset from South America (starting on Jan 1^st^, 2023) of FAO-confirmed H5N1 cases (https://microreact.org/project/vBkfwLENtQqwJ5G7YZfZDb-influenzasea-lionsacases) (32).

### Augur and individual ML tree display

TreeTime was employed to reconstruct ancestral sequences using a maximum likelihood approach and to map nucleotide mutations for each branch in the phylogenetic tree compared with the reference sequence A/Goose/Guangdong/1/96 (H5N1) (33).

## RESULTS

### HPAIV H5N1 outbreak in Uruguay

Out of 90 samples (72 pinnipeds and 18 seabirds) analyzed until 4 October, 29 pinnipeds and four terns tested positive for HPAIV H5N1 (Appendix Table 1). Samples for different locations with lower threshold cycles (Ct) were selected for genome sequencing (Table 1).

**TABLE 1:** Samples used in the study.

| Isolate (influenza type/host/country/strain ID/year) | Date | Location | Swab* | C<sub>t</sub><sup>†</sup> | Host | Symptoms/age/sex |
| --- | --- | --- | --- | --- | --- | --- |
| A/sea lion/Uruguay/P4_6923/2023 | 6 Sep 2023 | Maldonado (Punta del Este) | B | 23.61 | *Otaria flavescens* | Dead/adult |
| A/sea lion/Uruguay/P5_6923/2023 | 6 Sep 2023 | Canelones (El Pinar) | N | 29.24 | *Otaria flavescens* | Dead |
| A/sea lion/Uruguay/P6_6923/2023 | 6 Sep 2023 | Canelones (El Pinar) | O | 28.1 | *Otaria flavescens* | Neurological |
| **A/sea lion/Uruguay/P7_6923/2023** | 6 Sep<br>2023 | Canelones<br>(Médanos<br>de Solymar) | CSF | 28,1 | *Otaria*<br>*flavescens* | Dead |
| **A/fur seal/Uruguay/P8_8923/2023** | 8 Sep<br>2023 | Rocha (La<br>Paloma) | OP | 31,5 | *Arctocephalus*<br>*australis* | NA |
| **A/sea lion/Uruguay/P10_8923/2023** | 8 Sep<br>2023 | Rocha<br>(Cabo<br>Polonio) | OP/R | 25,1 | *Otaria*<br>*flavescens* | NA |
| **A/sea lion/Uruguay/P13_11923/2023** | 11 Sep<br>2023 | Canelones<br>(Atlántida) | N | 22.9 | *Otaria*<br>*flavescens* | Dead/M |
| **A/sea lion/Uruguay/P14_11923/2023** | 11 Sep<br>2023 | Canelones<br>(Parque del<br>Plata) | N/R | 23.5 | *Otaria*<br>*flavescens* | Dead/M |
| **A/sea lion/Uruguay/P15_14923/2023** | 14 Sep<br>2023 | Isla de<br>Flores | R | 29.3 | *Otaria*<br>*flavescens* | Male/juvenile |
| **A/tern/Uruguay/P16_14923/2023** | 14 Sep<br>2023 | Isla de<br>Flores | O/C | 20.2 | *Sterna*<br>*hirundinacea* | Dead/M/juvenile |
| **A/sea lion/Uruguay/P17_14923/2023** | 14 Sep<br>2023 | Isla de<br>Lobos | N | 25.5 | *Otaria*<br>*flavescens* | Dead |
| **A/sea lion/Uruguay/P18_14923/2023** | 14 Sep<br>2023 | Rocha<br>(Aguas<br>Dulce) | N | 23.8 | *Otaria*<br>*flavescens* | Dead/male |
| **A/tern/Uruguay/P23_41023/2023** | 4 Oct<br>2023 | Rocha<br>(Cabo<br>Polonio) | OP/C | 21,7 | *Sterna*<br>*hirundinacea* | Neurological |
| **A/tern/Uruguay/P24_41023/2023** | 4 Oct<br>2023 | Rocha<br>(Cabo<br>Polonio) | OP/C | 28,6 | *Sterna*<br>*hirundinacea* | Dead |
| **A/tern/Uruguay/P25_41023/2023** | 4 Oct<br>2023 | Rocha<br>(Cabo<br>Polonio) | OP | 24,5 | *Sterna*<br>*hirundinacea* | Neurological |
| **A/fur seal/Uruguay/P26_21023/2023** | 2 Oct<br>2023 | Rocha<br>(Cabo<br>Polonio) | N | 24,4 | *Arctocephalus*<br>*australis* | Dead/adult/F |

\*Types of samples: N: nasal; OP: oropharyngeal; C: cloacal; R: rectal; O: ocular; B; brain; CSF: cerebrospinal fluid.
<sup>†</sup>C<sub>t</sub> in the real-time reverse transcription PCR for the matrix gene.

### Genome variability and comparison

We obtained 16 viral genomes from ten sea lions (*Otaria flavescens*), two fur seals (*Arctocephalus australis*), and four terns (*Sterna hirundinacea*) (Table 1). Seven pinnipeds and three terns yielded complete viral genome sequences (Appendix Table 2). For the remaining samples (n=6), partial sequences comprising at least six genome segments were obtained.

The complete coding genome variability was low (distance>99.8%), and the segment sequences obtained showed the highest BLAST similarity, up to 100% similarity, with strains of the HPAI A/H5N1 2.3.4.4b lineage obtained from avian and mammalian hosts in South America.

### Phylogenetic analysis

Phylogenetic analyses were performed individually on the eight genome segments (Appendix Figure 1). Individual phylogenetic trees show similar relationships for the Uruguayan sequences; they were closely related and associated with Peruvian and Chilean strains. There is no evidence of reassortment events in the dataset, supporting the fact that all South American strains (n=162) were derived from strains with a single segment composition that successfully spread in South America. The 4:4 reassortant viruses, with segments derived from both North American (PB2, PB1, NP, and NS) and Eurasian (PA, HA, N, A, and M) lineages (12,13,20)

The South American dataset showed low sequence variability, with fewer informative phylogenetic sites per segment. We concatenated all coding sequences (CDS) for the genomes to augment the phylogenetic signal. The Uruguayan sequences from sea lions, fur seals, and terns were clustered in a monophyletic clade, denoted clade 5.1 (Figure 1A), with the highest posterior probability [PP] support of 1 (containing four unique nucleotide substitutions and one amino acid change). The 5.1 clade tMRCA was estimated in late June 2023 (95% High Probability Density [HPD] between early June and late July 2023) (Figure 1A and B – clade 5.1). This clade also clustered with high support (PP =1) to sequences from Peru (sea lions) and Chile (sea lions and a human case) in March 2023 (Figure 1A – clade 5). The clade 5 tMRCA dates to early January 2023, with 95% HPD between late December 2022 and late February 2023. A sanderling (*Calidris alba*) and a sea lion strain also sampled in Chile were basal to this clade 5 (PP = 1).

**Figure 1.**
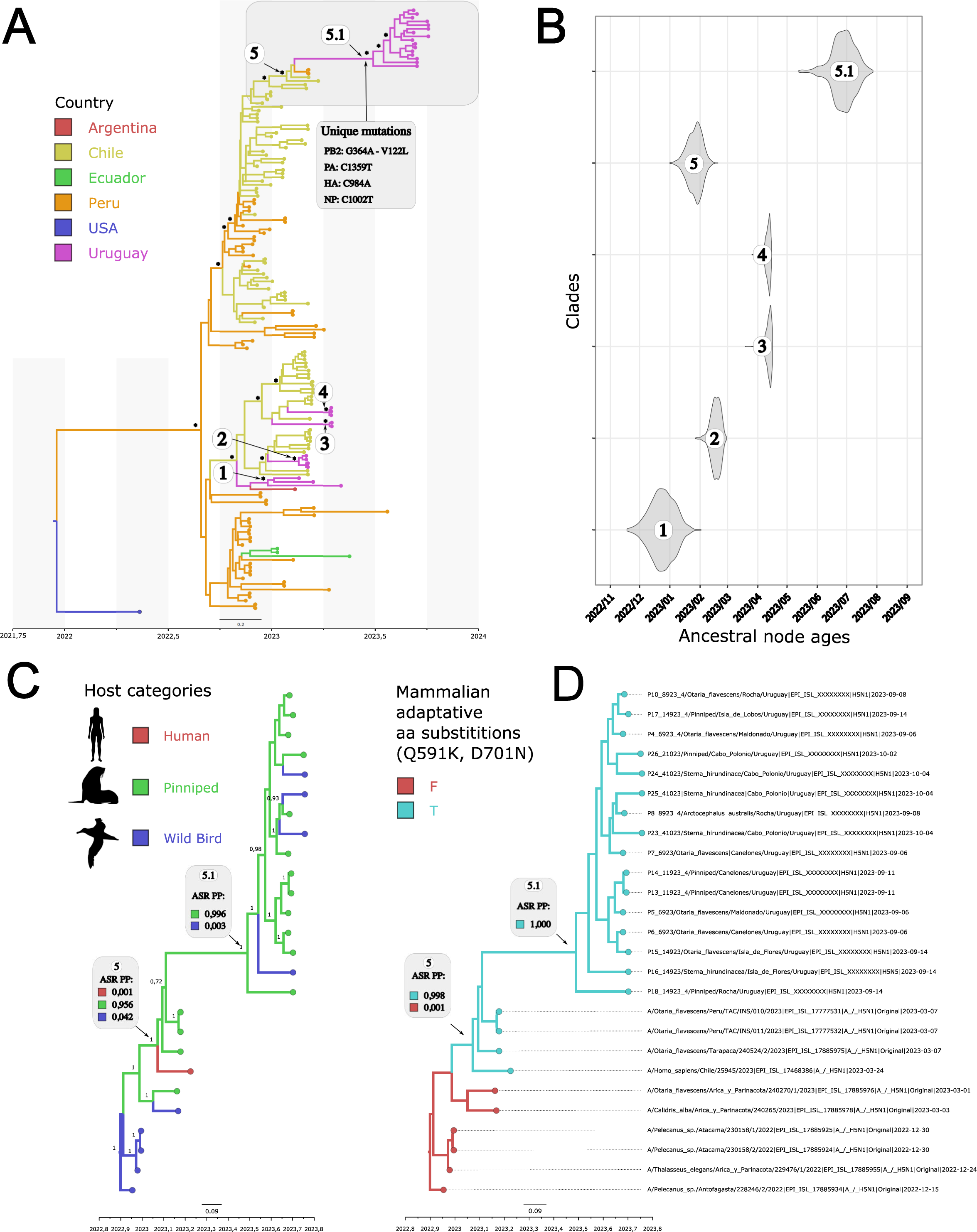
Bayesian phylogenetic reconstruction of concatenated segments, including timing estimates of the most recent common ancestors of specific Uruguayan clades and ancestral state reconstruction of countries, hosts, and presence/absence of two PB2 mutations known to increase the infection in mammalian cells. A – Discrete phylogeographic analysis of South American countries. Country colors denote specific countries, asterisks are posterior probability support above 90, and numbers indicate distinct Uruguayan clades. B – tMRCA of the six selected clades highlighted in part A. C – Discreate ancestral host reconstruction and associated ancestral state reconstruction posterior probabilities of each trait (ASR PP). D – Discrete ancestral reconstruction of PB2 mammal adaptation double mutation and associated ASR PP values.

The HPAIV of the February-May 2023 Uruguayan outbreaks, affecting wild birds and backyard poultry, are phylogenetically distant and did not form a monophyletic group with clade 5.1. They are associated with strains from Argentina, Peru, and Chile (Figure 1A and B– clades 1-4).

### Mammalian adaptive markers

Viruses from the Uruguayan pinnipeds and the four South American terns have the Q591K and D701N amino acid substitutions in PB2 (Figure 1, Appendix Table 3).

These substitutions occurred in only a few other South American strains: two strains from Peruvian sea lions, one from a Chilean sea lion, and one from a Chilean human case (Figure 1A–clade 5) (Appendix Table 4). All these viruses from Chile, Peru, and Uruguay, with 591K and 701N residues, form a monophyletic group (PP =1) with pinnipeds as the ancestral host (Figure 1C - clade 5) showing an ancestral state reconstruction posterior probability of 0.956.

Two other phylogenetically related but basal to clade 5, a sanderling, *Calidris alba* (EPI_ISL_17885978), and a sea lion (EPI_ISL_17885976) from Chile, have 701N but not 591K (Appendix Table 4). In this case, the ancestral host reconstruction showed a similar probability of 48 and 51% for avian and pinniped, respectively. The 701N and 591K residues in the PB2 segment were not observed for all remaining viral genomes analyzed.

Uruguayan samples belonging to clade 5.1 also depicted the substitutions PB2-V122I and PA-E237A (except for one virus), which were absent in any Chilean or Peruvian virus (Appendix Table 4).

### Genomic epidemiology

We reconstructed the phylogeographic dispersion of the H5N1 lineage in South America using discrete and continuous phylogeographic models. However, the phylogeographic reconstruction was impacted due to the almost complete lack of sequences from Southern Chile, Argentina, Bolivia, and Paraguay. The adjusted robust discrete phylogeographic analysis, considering the dataset’s large sampling gap, indicates that Uruguayan sequences from sea lions, fur seals, and terns likely originated from Chile (ASR PP = 0.9983).

Even though there are substantial sampling gaps in genomic surveillance in South America, there is consistent diagnostic data reported by FAO (https://empres-i.apps.fao.org/). Therefore, to visualize the case counts based on the larger animal categories (wild birds, pinnipeds, terrestrial mammals, and humans), we displayed all cases into two Microreact instances. One includes the concatenated maximum likelihood phylogenetic tree depicting it on the South American map and associated metadata (https://microreact.org/project/nSinj3LbMRme4gqussrTaU-influenzasea-lionurug). The second consists of all confirmed cases cataloged by FAO in South America after January 1^st^, 2023 (https://microreact.org/project/vBkfwLENtQqwJ5G7YZfZDb-influenzasea-lionsacases). Cases until 31 July 2023 showed an initial coastal and an internal spreading involving wild birds, backyard poultry, and outbreaks in commercial farms. After the first cases of pinnipeds in Peru, the virus spread to the South of the Pacific coast (Appendix Figure 2). From August 1^st^, registers appeared south of the Atlantic coast and spread northward, with fewer cases involving birds and case detections in the interior of South America (Appendix Figure 2).

### Intra-host variability of the sequenced genomes

Intra-host variation was observed within the virus segments in some samples, except for the PB1 segment. These variations were in the form of single nucleotide changes that sometimes affected the amino acid sequences (Appendix Table 5). Apart from a single nonsynonymous substitution in the M gene (H319N), minor variants were unique to each strain (i.e., they were not present in multiple sequences).

## DISCUSSION

From September to October 2023, HPAI subtype H5N1 caused the death of pinnipeds (*Otaria flavescens* and *Arctocephalus australis*) along 270 km of the Uruguayan coastline. Some terns (*Sterna hirundinacea*) were also found dead or with neurological symptoms a month after the first pinniped detections (Table 1, Appendix Table 1).

The estimated population size of the sea lion *Otaria flavescens* in Uruguay is almost 15,000 individuals (34), while the fur seal *A. australis* constitutes the biggest colony in South America, with approximately 300,000 individuals (35). The high mortality observed on the Uruguayan coast is comparable with that observed in Peru, Chile, and Argentina (11,12,36). Notably, sea lions were more affected than fur seals despite the latter’s larger population, suggesting a higher HPAI H5N1 susceptibility.

Some reports have described HPAIV in marine mammals outside South America. The reported species included harbor seals (*Phoca vitulina*) and gray seals (*Halichoerus grypus*) (16,37) in the northern hemisphere. Most of the cases were associated with clinical and pathological symptoms of pneumonia and were primarily detected from the respiratory samples (38–40). On the contrary, the latest reports in pinnipeds exhibited partial or extensive evidence of clinical neurological signs (11,12,16). In Uruguay, respiratory symptoms were found in one sea lion, and neurological symptoms in one sea lion and three terns (Table 1). It is possible that some animals showed respiratory or neurological symptoms before deceased, but most were found dead without any clinical signs recorded.

The pinniped outbreak was associated with HPAIV H5N1 infection in four terns. Despite active surveillance between June and October 2023, these are the first HPAV detections in wild birds in Uruguay. These seabirds inhabit coastal regions and share their habitat with marine mammals.

The avian influenza outbreak in pinnipeds and terns in Uruguay differs from that observed in wild birds and backyard poultry from February to May (13). These outbreaks belong to separate phylogenetic groups, harbor specific amino acid residues, and have non-overlapping tMRCA and ancestral hosts, supporting diverse geographic origins and a divergent transmission displacement (Figure 1A and B, Appendix Table 3). This indicates that Uruguay experienced two infection waves of HPAIV (H5N1) of the clade 2.3.4.4b during 2023. The first wave, affecting wild birds and backyard poultry, arrived in Uruguay within weeks of detection on the Pacific coast of Chile and expanded over four months (February to May 2023). Outbreaks appeared interspersed in a major clade with strains from Chile and Argentina, indicating the possibility of multiple introductions (13). The second monophyletic wave affects primarily pinnipeds and tern species, leading to an unusual mortality event in pinnipeds from September to October 2023. This latter behavior supports the idea that the second wave originates more recently from a common ancestor that entered the host population as a single event.

The second Uruguayan wave is characterized by strains harboring the 591K and 701N residues in the PB2 subunit of the RNA polymerase complex, a critical determinant of AIV host range and adaptation (41,42). The AIV polymerase performs poorly in mammalian cells, likely because its catalytic activity requires interaction with host proteins, such as importin-α and ANP32A, which differ between birds and mammals (42). Thus, to overcome host restriction of the polymerase, influenza virus strains in mammals usually acquire and select adaptive substitutions in PB2 (43). Studies of natural zoonoses and experimental passaging have identified mutations that adapt polymerases to mammalian hosts, such as the detected Q591K and D701N substitutions associated with increased polymerase activity in mammalian cells (44).

In the present analysis, PB2-591K and PB2-701N were restricted to the clade 5 comprised of three sea lions collected in March 2023 from Chile and Peru, a human case from Chile, and Uruguayan sea lions, fur seals, and terns (Figure 1A and https://microreact.org/project/nSinj3LbMRme4gqussrTaU-influenzasea-lionurug). The phylogenetic similarity, the 591K/701N residues, and the ancestral state reconstruction suggest that Uruguayan viruses originated from similar strains circulating in pinnipeds from the Chilean and Peruvian coasts (45,46). The expansion of this successful lineage might be related to the combined residues that enhance the geographic spreading ability of the virus through mammalian hosts. Data from current and previous influenza global outbreaks indicate that the simultaneous occurrence of both mutations is rarely observed in nature (20,47). The 591K and 701N residues in our study might have coevolved through adaptive selection, facilitating replication in mammalian hosts (47). Co-adaptation of PB2 mutations provides higher adaptability of AIVs in mammals and might contribute to crossing the interspecies barrier to mammals (48).

Our findings have two remarkable implications uncommon for HPAIV epidemiology: the plausibility of mammal-to-mammal and mammal-to-bird transmissions.

Mammal-to-mammal transmission of HPAIV H5N1 is rare but has been proposed in captive tigers and experiments with cats (49,50). However, interspecific transmission from experimentally infected cats to naïve domestic dogs did not occur (51). Recent research also suggested mammal-to-mammal infection of the clade 2.3.4.4b in fur farms (14,52).

Wild pinniped populations have peculiar characteristics like those observed in mammals kept in captivity. These highly social animals frequently interact through physical contact, fights, mate competition, vocalizations such as spitting and hissing, and have constant contact with the feces and urine of other colony animals. This interplay and the high-density populations may facilitate the transmission of HPAIV and serve as a model for studying mammal-to-mammal transmission in natural environments.

Influenza in pinnipeds spread throughout the coastline of Peru, Chile, Argentina, Uruguay, and recently Brazil (22,36). This spread by proximity along the southern coast of South America, from the Pacific to the Atlantic Ocean (Appendix Figure 2), is consistent with the virus being transmitted mainly by pinnipeds. A primary mammal-to-mammal transmission also explains why the Uruguayan and Chilean strains of the virus are closely related, even though they were collected six months apart, and that many sea lions are currently affected simultaneously. We proposed that only the lineage harboring PB2-591K/701N mammalian adaptive residues expanded to Argentina, Uruguay, and Brazil. Unlike what has been observed in Chile and Peru, this expansion on the Atlantic coast was not preceded and concurrent with massive bird cases despite intensive surveillance (Appendix Figure 2) (36).

The phylogenetic results and the 591K and 701N mammalian markers in tern viruses suggested that they have mammalian origin and were directly acquired from sea lions and fur seals on the Uruguayan coasts. Pinnipeds spend time ashore in sites typically used by seabirds for roosting, facilitating cross-species transmission of AIV between these species.

Mammalian adaptive markers are linked with the passage of the virus through a mammalian host, and it is unlike that they developed *de novo* in bird hosts. Moreover, as described in Chile, we have not observed intra-host variability (minor single nucleotide variants) in the PB2-591/701 codons in marine mammals or avian genomes that support an ongoing change in mammalian adaptive residues (Appendix Table 5) (20). The ability of the virus to retain acquired genetic variability during the mammal passage is of significant concern since these viruses could be transmitted to other mammals, including humans, directly and potentially indirectly by affecting wild birds or poultry populations. Although these mammalian adaptations caused the death of some individuals (Table 1), there have not been massive death events registered in seabirds in the Uruguayan coastal region, suggesting that the virus has a lower spreading ability among birds.

The main limitation of our studies is the knowledge gap in the sequences available from some South American countries, particularly Argentina, that affects the performance of phylogeographic inferences. Another concern is the lack of clinical data and the unavailability of tissue and blood samples to further the analysis.

Our study introduces a new perspective on influenza transmission in marine mammals in South America. It also offers an additional framework for understanding how avian influenza spreads and impacts bird and mammalian hosts. The challenge of controlling avian influenza is amplified by its spread across hosts adapted to air, water, and water-land interfaces. Viruses capable of infecting and replicating in pinnipeds may be more adapted to mammalian than avian hosts (20). In addition to serving as a spillover host, other influenza subtypes could be endemic in Uruguayan pinnipeds (53), providing genetic reassortment opportunities. Lastly, the AIV interspecies transmission should be closely monitored through continued surveillance and containment measures to restrict transmission.

## Acknowledge

We want to express our gratitude to Valeria Gayo from DILAVE for effectively coordinating the research activities in this study.

We thank the Armada Nacional del Uruguay for providing transportation via naval vessels to the Ministry of Environment’s researchers.

## Institutional Review Board Statement

Ethical approval was waived because the study was conducted as part of public health surveillance by official authorities in response to avian flu outbreaks.

## Appendix

**Appendix Table 1.** Highly pathogenic **a**vian influenza H5N1 virus-positive cases in Uruguayan samples.

**Appendix Table 2.** Coverage of the different highly pathogenic avian influenza virus H5N1 segments from Uruguayan strains.

**Appendix Table 3.** Amino acid differences between highly pathogenic avian influenza H5N1 virus invasions in Uruguay at different times. Wave I (Feb-May 2023): wild birds and backyard poultry. Wave II (Sep-October 2023): pinnipeds and seabirds.

**Appendix Table 4.** Amino acid changes between Uruguayan (wave II) and Chilean/Peruvian highly pathogenic avian influenza H5N1 virus strains.

**Appendix Table 5.** Single nucleotide intra-host variant in Uruguayan strains of the highly pathogenic avian influenza H5N1 virus.

**Appendix Figure 1.** Phylogenetic analysis of the highly pathogenic avian influenza H5N1 virus coding regions from South America. Uruguayan strains are highlighted in light brown (Wave I) and light blue (Wave II). Hosts and mammalian adaptive residues (PB2-591Q/701N) are depicted. A) Maximum-likelihood tree based on segment 1 (PB2); B) Maximum-likelihood tree based on segment 2 (PB1); C) Maximum-likelihood tree based on segment 3 (PA); D) Maximum-likelihood tree based on segment 4 (HA); E) Maximum-likelihood tree based on the segment 5 (NP); F) Maximum-likelihood tree based on the segment 6 (NA); G) Maximum-likelihood tree based on the segment 7 (M); H) Maximum-likelihood tree based on the segment 8 (NS).

**Appendix Figure 2:** Variation in the distribution of the highly pathogenic avian influenza H5N1 avian influenza virus in South America (Microreact instance including all confirmed cases cataloged by FAO in South America after January 1^st^, 2023). A) Cases until 31 July 2023. Registers showed an initial coastal and an internal spreading involving wild birds, backyard poultry, and outbreaks in commercial farms. B) Cases from 1^st^ August to October 2023. Registers appeared south of the Atlantic coast and spread northward, with fewer cases involving birds and case detections in the interior of South America.

