## Appendix for "Highly pathogenic avian influenza H5N1 virus infections in pinnipeds and seabirds in Uruguay: a paradigm shift to virus transmission in South America"

Tables and Figures

Tomás et al. 2023

**Table legends**

**Appendix Table 1.** Highly pathogenic **a**vian influenza H5N1 virus-positive cases in Uruguayan samples.

**Appendix Table 2.** Coverage of the different highly pathogenic avian influenza virus H5N1 segments from Uruguayan strains.

**Appendix Table 3.** Amino acid differences between highly pathogenic avian influenza H5N1 virus invasions in Uruguay at different times. Wave I (Feb-May 2023): wild birds and backyard poultry. Wave II (Sep-October 2023): pinnipeds and seabirds.

**Appendix Table 4.** Amino acid changes between Uruguayan (wave II) and Chilean/Peruvian highly pathogenic avian influenza H5N1 virus strains.

**Appendix Table 5.** Single nucleotide intra-host variant in Uruguayan strains of the highly pathogenic avian influenza H5N1 virus.

**Appendix Table 1.** Highly pathogenic **a**vian influenza H5N1 virus-positive cases in Uruguayan samples.

| Positive cases | Date* | Department | Location | Host class | Host species |
|---|---|---|---|---|---|
| 1 | 2023 Sep 4 | Montevideo | Playa del Cerro | Mammal | *Otaria flavescens* |
| 2 | 2023 Sep 6 | Maldonado | Punta del Este | Mammal | *Arctocephalus australis* |
| 3 | 2023 Sep 6 | Maldonado | Punta del Este | Mammal | *Otaria flavescens* |
| 4 | 2023 Sep 6 | Maldonado | Punta del Este | Mammal | *Otaria flavescens* |
| 5 | 2023 Sep 6 | Canelones | El Pinar | Mammal | Otariid |
| 6 | 2023 Sep 6 | Canelones | El Pinar | Mammal | Otariid |
| 7 | 2023 Sep 6 | Canelones | Lomas de Solymar | Mammal | Otariid |
| 8 | 2023 Sep 8 | Rocha | La Paloma | Mammal | *Arctocephalus australis* |
| 9 | 2023 Sep 8 | Rocha | Cabo Polonio | Mammal | *Otaria flavescens* |
| 10 | 2023 Sep 8 | Rocha | Cabo Polonio | Mammal | *Otaria flavescens* |
| 11 | 2023 Sep 11 | Canelones | Shangrilá | Mammal | Pinniped |
| 12 | 2023 Sep 11 | Canelones | Atlántida | Mammal | *Otaria flavescens* |
| 13 | 2023 Sep 11 | Canelones | Parque del Plata | Mammal | *Otaria flavescens* |
| 14 | 2023 Sep 14 | Canelones/Montevideo | Isla de Flores | Mammal | *Otaria flavescens* |
| 15 | 2023 Sep 14 | Canelones/Montevideo | Isla de Flores | Bird | *Sterna hirundinacea* |
| 16 | 2023 Sep 14 | Maldonado | Isla de Lobos | Mammal | Pinniped |
| 17 | 2023 Sep 14 | Maldonado | Isla de Lobos | Mammal | *Otaria flavescens* |
| 18 | 2023 Sep 14 | Rocha | Cabo Polonio | Mammal | Pinniped |
| 19 | 2023 Sep 14 | Rocha | Cabo Polonio | Mammal | Pinniped |
| 20 | 2023 Sep 14 | Rocha | Cabo Polonio | Mammal | Pinniped |
| 21 | 2023 Sep 14 | Rocha | Cabo Polonio | Mammal | Pinniped |
| 22 | 2023 Sep 14 | Rocha | Aguas Dulces | Mammal | *Otaria flavescens* |
| 23 | 2023 Oct 2 | Rocha | Cabo Polonio | Mammal | Pinniped |
| 24 | 2023 Oct 2 | Rocha | Cabo Polonio | Mammal | *Arctocephalus australis* |
| 25 | 2023 Oct 2 | Rocha | Cabo Polonio | Mammal | Pinniped |
| 26 | 2023 Oct 2 | Rocha | Cabo Polonio | Mammal | Pinniped |
| 27 | 2023 Oct 2 | Rocha | Cabo Polonio | Mammal | Pinniped |
| 28 | 2023 Oct 2 | Rocha | Cabo Polonio | Mammal | Pinniped |
| 29 | 2023 Oct 2 | Rocha | Cabo Polonio | Mammal | Pinniped |
| 30 | 2023 Oct 2 | Rocha | Cabo Polonio | Mammal | Pinniped |
| 31 | 2023 Oct 4 | Rocha | Cabo Polonio | Bird | *Sterna hirundinacea* |
| 32 | 2023 Oct 4 | Rocha | Cabo Polonio | Bird | *Sterna hirundinacea* |
| 33 | 2023 Oct 4 | Rocha | Cabo Polonio | Bird | *Sterna hirundinacea* |

*Date of sample received.

**Appendix Table 2.** Coverage of the different highly pathogenic avian influenza virus H5N1 segments from Uruguayan strains.

| Sample | Mean depth of coverage (Std Dev) | |
|---|---|---|
| P4_6923_*Otaria flavescens* | | |
| PB2 gene | 187.7 (31.4)[*1] | |
| PB1 gene | 796.5 (619.8)[*1] | |
| PA gene | 912.6 (124.0)[*1] | |
| HA gene | 149.2 (149.8)[*1] | |
| NP gene | 536.2 (66.2)[*1] | |
| NA gene | 78.8 (91.5)[*1] | |
| M gene | 361.2 (69.4)[*1] | |
| NS gene | 272.5 (38.2)[*1] | |
| P5_6923_*Otaria flavescens* | | |
| PB2 gene | | 16.6 (5.8)[*2] |
| PB1 gene | | NC[†] |
| PA gene | | 848.8 (156.4)[*2] |
| HA gene | | NC |
| NP gene | | 721.9 (96.9)[*2] |
| NA gene | | 545.1 (146.5)[*2] |
| M gene | | 1161.1(187.3)[*2] |
| NS gene | | 4004.8 (404.6)[*2] |
| P6_6923_*Otaria flavescens* | | |
| PB2 gene | | 315.1 (69.9)[*2] |
| PB1 gene | | NC |
| PA gene | | 1017.2 (185.8)[*2] |
| HA gene | | 174.6 (232.8)[*2] |
| NP gene | | 622.8 (106.7)[*2] |
| NA gene | | 836.2 (194.4)[*2] |
| M gene | | 2208.8 (368.5)[*2] |
| NS gene | | 2673.1 (268.9)[*2] |
| P7_6923_*Otaria flavescens* | | |
| PB2 gene | | 121.6 (85.7)[*2] |
| PB1 gene | | NC |
| PA gene | | 1237.7 (759.1)[*2] |
| HA gene | | 257.6 (303.5)[*2] |
| NP gene | | 339.0 (55.6)[*2] |
| NA gene | | 481.0 (124.1)[*2] |
| M gene | | 503.9 (84.5)[*2] |
| NS gene | | 2584.1 (258.7)[*2] |
| P8_8923_*Arctocephalus australis* | | |
| PB2 gene | 9.6 (3.4)[*1] | |
| PB1 gene | 59.4 (12.4)[*1] | |
| PA gene | 57.1 (10.9)[*1] | |
| HA gene | 22.5 (5.9)[*1] | |
| NP gene | 501.2 (81.5)[*1] | |

| | | |
|---|---|---|
| NA gene | 44.0 (10.5)*1 | |
| M gene | 5794.1 (1100.4)*1 | |
| NS gene | 2882.7 (324.4)*1 | |
| P10_8923_*Otaria flavescens* | | |
| PB2 gene | 925.3 (128.7)*1 | 87.2 (28.4)*2 |
| PB1 gene | 1215.6 (370.5)*1 | 99.4 (34.8)*2 |
| PA gene | 1775.1 (212.4)*1 | 247.4 (33.8)*2 |
| HA gene | 5513.1 (3071.7)*1 | 481.7 (6533.9)*2 |
| NP gene | 7812.3 (794.9)*1 | 5371.4 (620.5)*2 |
| NA gene | 1898.5 (1139.5)*1 | 2872.2 (647.4)*2 |
| M gene | 32681.9 (5550.4)*1 | 1743.4 (367.7)*2 |
| NS gene | 9520.4 (1332.5)*1 | 2684.6 (560.1)*2 |
| P13_11923_*Otaria flavescens* | | |
| PB2 gene | 434.0 (80.8)*1 | |
| PB1 gene | 798.4 (246.0)*1 | |
| PA gene | 964.2 (170.0)*1 | |
| HA gene | 917.4 (524.5)*1 | |
| NP gene | 1939.9 (228.1)*1 | |
| NA gene | 298.6 (254.2)*1 | |
| M gene | 3532.0 (612.2)*1 | |
| NS gene | 2435.1 (387.1)*1 | |
| P14_11923_*Otaria flavescens* | | |
| PB2 gene | 35.1 (6.6)*1 | 2292.7 (305.0)*2 |
| PB1 gene | 66.4 (17.0)*1 | 6.4 (3.6)*2 |
| PA gene | 55.4 (12.8)*1 | 9412.5 (1039.2)*2 |
| HA gene | 58.5 (25.3)*1 | 146.0 (161.3)*2 |
| NP gene | 153.5 (20.1)*1 | 3747.5 (486.5)*2 |
| NA gene | 8.0 (5.3)*1 | 12336.0 (2400.7)*2 |
| M gene | 202.5 (38.4)*1 | 37618.0 (8382.3)*2 |
| NS gene | 71.1 (12.7)*1 | 6608.4 (1502.7)*2 |
| P15_14923_*Otaria flavescens* | | |
| PB2 gene | | 341.4 (340.7)*2 |
| PB1 gene | | NC |
| PA gene | | NC |
| HA gene | | 225.4 (285.4)*2 |
| NP gene | | 1812.2 (336.6)*2 |
| NA gene | | 546.5 (137.7)*2 |
| M gene | | 520.8 (88.4)*2 |
| NS gene | | 2890.1 (343.0)*2 |
| P16_14923_ *Sterna hirundinacea* | | |
| PB2 gene | 310.9 (82.7)*1 | 404.7 (149.6)*2 |
| PB1 gene | 647.3 (335.5)*1 | 48.7 (20.0)*2 |
| PA gene | 505.9 (93.4)*1 | 742.6 (123.9)*2 |
| HA gene | 3801.4 (2494.5)*1 | 15274.3 (8070.0)*2 |

| | | | |
|---|---|---|---|
| NP gene | 5165.8 (784.5)[*1] | | 14723.5 (1661.4)[*2] |
| NA gene | 1370.1 (858.2)[*1] | | 3438.5 (535.3)[*2] |
| M gene | 34512.2 (6018.9)[*1] | | 2215.8 (454.9)[*2] |
| NS gene | 13722.1 (2212.0)[*1] | | 1433.6 (286.1)[*2] |
| P17_14923_*Otaria flavescens* | | | |
| PB2 gene | 5.0 (2.3)[*1] | | |
| PB1 gene | 6.4 (3.3)[*1] | | |
| PA gene | 6.7 (2.7)[*1] | | |
| HA gene | 9.6 (3.5)[*1] | | |
| NP gene | 24.7 (7.2)[*1] | | |
| NA gene | 2.0 (1.2)[*1] | | |
| M gene | 36.0 (8.7)[*1] | | |
| NS gene | 23.4 (4.0)[*1] | | |
| P18_14923_*Otaria flavescens* | | | |
| PB2 gene | 20.4 (5.4)[*1] | | 2516.9 (354.9)[*2] |
| PB1 gene | 18.5 (6.5)[*1] | | 10.5 (4.3)[*2] |
| PA gene | 25.2 (4.6)[*1] | | 7948.3 (1024.8)[*2] |
| HA gene | 47.4 (14.7)[*1] | | 497.0 (546.1)[*2] |
| NP gene | 86.0 (16.1)[*1] | | 6739.5 (1064.4)[*2] |
| NA gene | 19.9 (4.8)[*1] | | 15310.7 (3061.2)[*2] |
| M gene | 231.9 (41.9)[*1] | | 29767.8 (6517.3)[*2] |
| NS gene | 161.1 (31.0)[*1] | | 14944.5 (3311.9)[*2] |
| P23_41023_*Sterna hirundinacea* | | | |
| PB2 gene | | 745.0 (452.2)[*12] | |
| PB1 gene | | 32.7 (16.5)[*12] | |
| PA gene | | 1358.3 (397.9)[*12] | |
| HA gene | | 1256.8 (996.9)[*12] | |
| NP gene | | 2294.1 (371.8)[*12] | |
| NA gene | | 2226.3 (665.5)[*12] | |
| M gene | | 3414.6 (538.6)[*12] | |
| NS gene | | 10383.5 (904.9)[*12] | |
| P24_41023_*Sterna hirundinacea* | | | |
| PB2 gene | | 86.3 (17.8)[*12] | |
| PB1 gene | | NC | |
| PA gene | | 130.2 (28.1)[*12] | |
| HA gene | | 115.5 (40.6)[*12] | |
| NP gene | | 1493.5 (223.9)[*12] | |
| NA gene | | NC | |
| M gene | | 287.1 (59.5)[*12] | |
| NS gene | | 3666.9 (378.7)[*12] | |
| P25_41023_*Sterna hirundinacea* | | | |
| PB2 gene | | 2192.4 (796.3)[*12] | |
| PB1 gene | | 27.5 (55.5)[*12] | |
| PA gene | | 2204.2 (412.4)[*12] | |

| | |
|---|---|
| HA gene | 333.7 (94.2)*[12] |
| NP gene | 907.5 (128.7)*[12] |
| NA gene | 94.4 (133.6)*[12] |
| M gene | 6513.8 (1235.3)*[12] |
| NS gene | 11985.2 (1131.6)*[12] |
| P26_21023_ *Arctocephalus australis* | |
| PB2 gene | 888.5 (1341.7)*[12] |
| PB1 gene | NC |
| PA gene | 1275.7 (1357.1)*[12] |
| HA gene | 716.3 (888.0)*[12] |
| NP gene | 1079.6 (177.0)*[12] |
| NA gene | 999.4 (261.4)*[12] |
| M gene | 2412.7 (360.2)*[12] |
| NS gene | 8446.2 (756.8)*[12] |

*[1]Library preparation from amplicons, according to Zhou et al. (2009).

*[2]Library preparation from amplicons, according to Hoffmann et al. (2001).

*[12]Library preparation from both PCR methods.
[†]Non-coverage.

**Appendix Table 3**. Amino acid differences between highly pathogenic avian influenza H5N1 virus invasions in Uruguay at different times. Wave I (Feb-May 2023): wild birds and backyard poultry. Wave II (Sep-October 2023): pinnipeds and seabirds.

| | | PB2 | | | | PB1 | | PA | | | | | NP | NS1 | | | |
|---|---|---|---|---|---|---|---|---|---|---|---|---|---|---|---|---|---|
| | Strain | 122 | 154 | **591*** | **701*** | 515 | 646 | 20 | 57 | 86 | 237 | 548 | 119 | 21 | 26 | 53 | 226 |
| Wave I | EPI_ISL_18310958 | V | - | **Q** | **D** | S | I | A | R | M | E | M | I | R | E | D | I |
| | EPI_ISL_18310942 | · | F | **·** | **·** | · | · | · | · | · | · | · | · | · | · | · | · |
| | EPI_ISL_18310967 | · | · | **·** | **·** | · | · | · | · | · | · | · | · | · | · | · | · |
| | EPI_ISL_18310966 | · | · | **·** | **·** | · | · | · | · | · | · | · | · | · | · | · | · |
| | EPI_ISL_18310965 | · | · | **·** | **·** | · | · | · | · | · | · | · | · | · | · | · | · |
| | EPI_ISL_18310957 | · | · | **·** | **·** | · | · | · | · | · | · | · | · | · | · | · | · |
| | EPI_ISL_18310964 | · | · | **·** | **·** | · | · | · | · | · | · | · | · | · | · | · | · |
| | EPI_ISL_18310960 | · | · | **·** | **·** | · | · | · | · | · | · | · | · | · | · | · | · |
| | EPI_ISL_18310959 | · | · | **·** | **·** | · | · | · | · | · | · | · | · | · | · | · | · |
| | EPI_ISL_18310963 | · | · | **·** | **·** | · | · | · | · | · | · | · | · | · | · | · | · |
| | EPI_ISL_18310962 | · | · | **·** | **·** | · | · | · | · | · | · | · | · | · | · | · | · |
| | EPI_ISL_18310961 | · | · | **·** | **·** | · | · | · | · | · | · | · | · | · | · | · | · |
| Wave II | P4_6923 | I | L | **K** | **N** | A | M | T | Q | I | A | I | T | Q | K | G | T |
| | P5_6923 | I | L | **K** | **N** | - | - | T | Q | I | A | I | T | Q | K | G | T |
| | P6_6923 | I | L | **K** | **N** | - | - | T | Q | I | A | I | T | Q | K | G | T |
| | P7_6923 | I | L | **K** | **N** | A | M | T | Q | I | A | I | T | Q | K | G | T |
| | P8_8923 | I | L | **K** | **N** | A | M | T | Q | I | A | I | T | Q | K | G | T |
| | P10_8923 | I | L | **K** | **N** | A | M | T | Q | I | A | I | T | Q | K | G | T |
| | P13_11923 | I | L | **K** | **N** | A | M | T | Q | I | A | I | T | Q | K | G | T |
| | P14_11923 | I | L | **K** | **N** | A | M | T | Q | I | A | I | T | Q | K | G | T |
| | P15_14923 | I | L | **K** | **N** | A | M | T | - | - | - | - | T | Q | K | G | T |
| | P16_14923 | I | L | **K** | **N** | A | M | T | Q | I | A | I | T | Q | K | C | T |
| | P17_14923 | I | L | **K** | **N** | A | M | T | Q | I | A | I | T | Q | K | G | T |
| | P18_14923 | I | L | **K** | **N** | A | M | T | Q | I | · | I | T | Q | K | G | T |
| | P23_41023 | I | L | **K** | **N** | A | M | T | Q | I | A | I | T | Q | K | G | T |
| | P24_41023 | I | L | **K** | **N** | A | M | T | Q | I | A | I | T | Q | K | G | T |
| | P25_41023 | I | L | **K** | **N** | A | M | T | Q | I | A | I | T | Q | K | G | T |
| | P26_21023 | I | L | **K** | **N** | A | M | T | Q | I | A | I | T | Q | K | G | T |

*Markers associated with avian influenza virus adaptation to mammals.

**Appendix Table 4**. Amino acid changes between Uruguayan (wave II) and Chilean/Peruvian highly pathogenic avian influenza H5N1 virus strains.

| | | PB2 | | | PB1 | PA | | | | | NP | NS1 | | | |
|---|---|---|---|---|---|---|---|---|---|---|---|---|---|---|---|
| | Strain | 122 | **591*** | **701*** | 515 | 20 | 57 | 86 | 237 | 548 | 119 | 21 | 26 | 53 | 226 |
| Chile/Peru | EPI_ISL_18054503/dolphin/Peru | V | **Q** | **D** | S | A | R | M | E | M | I | R | E | D | I |
| | EPI_ISL_18054502/sea lion/Peru | · | · | · | · | · | · | I | · | · | · | · | · | · | · |
| | EPI_ISL_17805999/lion/Peru | · | · | · | · | · | · | · | · | · | · | · | · | · | · |
| | EPI_ISL_18054509/sea lion/Peru | · | · | · | · | · | Q | · | · | · | · | · | · | · | · |
| | EPI_ISL_17885978/sanderling/Chile | · | · | **N** | A | · | Q | · | · | · | T | · | K | G | T |
| | EPI_ISL_17885976/sea lion/Chile | · | · | **N** | A | · | Q | · | · | · | T | · | K | G | T |
| | EPI_ISL_17468386/human/Chile | · | **K** | **N** | · | T | Q | I | · | I | T | Q | K | G | T |
| | EPI_ISL_17777531/sea lion/Peru | · | **K** | **N** | A | · | Q | I | · | I | T | Q | K | G | T |
| | EPI_ISL_17777532/sea lion/Peru | · | **K** | **N** | A | · | Q | I | · | I | T | Q | K | G | T |
| | EPI_ISL_17885975/sea lion/Chile | · | **K** | **N** | · | · | Q | I | · | I | T | Q | K | G | T |
| Wave 2 | P4_6923 | I | **K** | **N** | A | T | Q | I | A | I | T | Q | K | G | T |
| | P5_6923 | I | **K** | **N** | - | T | Q | I | A | I | T | Q | K | G | T |
| | P6_6923 | I | **K** | **N** | - | T | Q | I | A | I | T | Q | K | G | T |
| | P7_6923 | I | **K** | **N** | A | T | Q | I | A | I | T | Q | K | G | T |
| | P8_8923 | I | **K** | **N** | A | T | Q | I | A | I | T | Q | K | G | T |
| | P10_8923 | I | **K** | **N** | A | T | Q | I | A | I | T | Q | K | G | T |
| | P13_11923 | I | **K** | **N** | A | T | Q | I | A | I | T | Q | K | G | T |
| | P14_11923 | I | **K** | **N** | A | T | Q | I | A | I | T | Q | K | G | T |
| | P15_14923 | I | **K** | **N** | A | T | - | - | - | - | T | Q | K | G | T |
| | P16_14923 | I | **K** | **N** | A | T | Q | I | A | I | T | Q | K | C | T |
| | P17_14923 | I | **K** | **N** | A | T | Q | I | A | I | T | Q | K | G | T |
| | P18_14923 | I | **K** | **N** | A | T | Q | I | · | I | T | Q | K | G | T |
| | P23_41023 | I | **K** | **N** | A | T | Q | I | A | I | T | Q | K | G | T |
| | P24_41023 | I | **K** | **N** | A | T | Q | I | A | I | T | Q | K | G | T |
| | P25_41023 | I | **K** | **N** | A | T | Q | I | A | I | T | Q | K | G | T |
| | P26_21023 | I | **K** | **N** | A | T | Q | I | A | I | T | Q | K | G | T |

*Markers associated with avian influenza virus adaptation to mammals.

**Appendix Table 5.** Single nucleotide intra-host variant in Uruguayan strains of the highly pathogenic avian influenza H5N1 virus.

| Sample/codon position | Change | Frequency (%) | Coverage | Polymorphism type |
|---|---|---|---|---|
| P5_6923_*Otaria flavescens* | | | | |
| PA_147 codon | A > G | 88 - 11.4 | 852 | synonymous |
| NA_73 codon | T > A | 88.4 - 11.6 | 859 | synonymous |
| M_19 codon | C > T | 88.7 - 10.9 | 1241 | synonymous |
| M_158 codon | G > A | 39.1 - 60.5 | 958 | synonymous |
| NS1_66 codon | G > A | 89.5 - 10.4 | 4460 | synonymous |
| P6_6923_*Otaria flavescens* | | | | |
| PB2_67 codon | A > G | 87.2 - 12.8 | 290 | I > V |
| NP_50 codon | A > G | 62.9 - 36.8 | 587 | S > G |
| P7_6923_*Otaria flavescens* | | | | |

| | | | | |
|---|---|---|---|---|
| NP_166 codon | G > T | 78.8 - 20.9 | 392 | synonymous |
| P8_8923_*Arctocephalus australis* | | | | |
| M_82 codon | T > A | 87.2 - 12.8 | 6475 | N > K |
| P13_11923_*Otaria flavescens* | | | | |
| PA_221 codon | G > A | 88.9 - 11.1 | 835 | synonymous |
| P15_14923_*Otaria flavescens* | | | | |
| NP_104 codon | C > A | 85.4 - 14.4 | 3512 | Q > K |
| M_319 codon | C > A | 55.4 - 44.4 | 597 | H > N |
| NS1_36 codon | C > A | 87.8 - 12.1 | 3006 | synonymous |
| NS1_122 codon | C > A | 85.6 - 14.4 | 2566 | synonymous |
| P23_41023_*Sterna hirundinacea* | | | | |
| NP_375 codon | G > A | 82.3 - 17.5 | 2246 | D > N |
| NA_169 codon | C > T | 89.1 - 10.7 | 1603 | P > S |
| P25_41023_*Sterna hirundinacea* | | | | |
| PB2_136 codon | G > A | 76 - 23.6 | 1507 | R > K |
| PB2_545 codon | G > A | 75.6 - 24.3 | 1682 | synonymous |
| HA_517 codon | A > G | 78.3 - 21.5 | 447 | synonymous |
| NP_223 codon | T > C | 85 - 14.7 | 822 | synonymous |
| P26_21023_ *Arctocephalus australis* | | | | |
| PB2_13 codon | G > A | 81.2 - 18.8 | 6142 | synonymous |
| PA_658 codon | T > G | 86.8 - 13.1 | 2761 | F > L |
| NP_422 codon | A > G | 82.1 - 17.9 | 944 | synonymous |
| M_186 codon | C > T | 70.9 – 29 | 2123 | synonymous |
| M_237 codon | G > A | 84.8 - 15.1 | 1776 | synonymous |
| M_319 codon | C > A | 87.7 - 12.2 | 2591 | H > N |

#### Figure legends

**Appendix Figure 1**. Phylogenetic analysis of the highly pathogenic avian influenza H5N1 virus coding regions from South America. Uruguayan strains are highlighted in light brown (Wave I) and light blue (Wave II). Hosts and mammalian adaptive residues (PB2-591Q/701N) are depicted. A) Maximum-likelihood tree based on segment 1 (PB2); B) Maximum-likelihood tree based on segment 2 (PB1); C) Maximum-likelihood tree based on segment 3 (PA); D) Maximum-likelihood tree based on segment 4 (HA); E) Maximum-likelihood tree based on the segment 5 (NP); F) Maximum-likelihood tree based on the segment 6 (NA); G) Maximum-likelihood tree based on the segment 7 (M); H) Maximum-likelihood tree based on the segment 8 (NS).

**Appendix Figure 2**: Variation in the distribution of the highly pathogenic avian influenza H5N1 avian influenza virus in South America (Microreact instance including all confirmed cases cataloged by FAO in South America after January 1$^{st}$, 2023). A) Cases until 31 July 2023. Registers showed an initial coastal and an internal spreading involving wild birds, backyard poultry, and outbreaks in commercial farms. B) Cases from 1$^{st}$ August to October 2023. Registers appeared south of the Atlantic coast and spread northward, with fewer cases involving birds and case detections in the interior of South America.

**Appendix Figure 1**. Phylogenetic analysis of the highly pathogenic avian influenza H5N1 virus coding regions from South America. Uruguayan strains are highlighted in light brown (Wave I) and light blue (Wave II). Hosts and mammalian adaptive residues (PB2-591Q/701N) are depicted. A) Maximum-likelihood tree based on segment 1 (PB2); B) Maximum-likelihood tree based on segment 2 (PB1); C) Maximum-likelihood tree based on segment 3 (PA); D) Maximum-likelihood tree based on segment 4 (HA); E) Maximum-likelihood tree based on the segment 5 (NP); F) Maximum-likelihood tree based on the segment 6 (NA); G) Maximum-likelihood tree based on the segment 7 (M); H) Maximum-likelihood tree based on the segment 8 (NS).

Appendix Figure 1A (segment 1 PB2 gene)

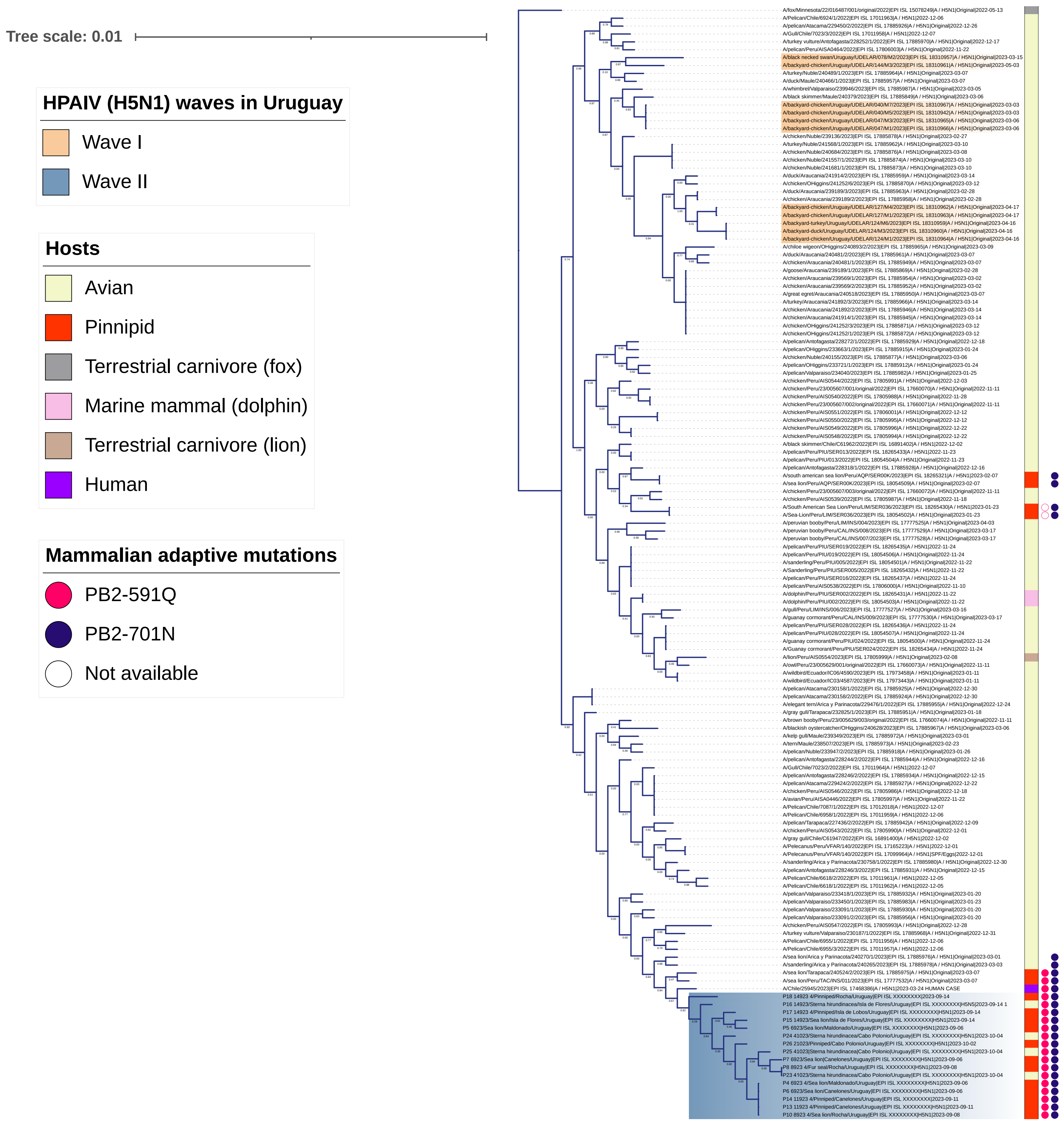

Appendix Figure 1B (segment 2 PB1 gene)

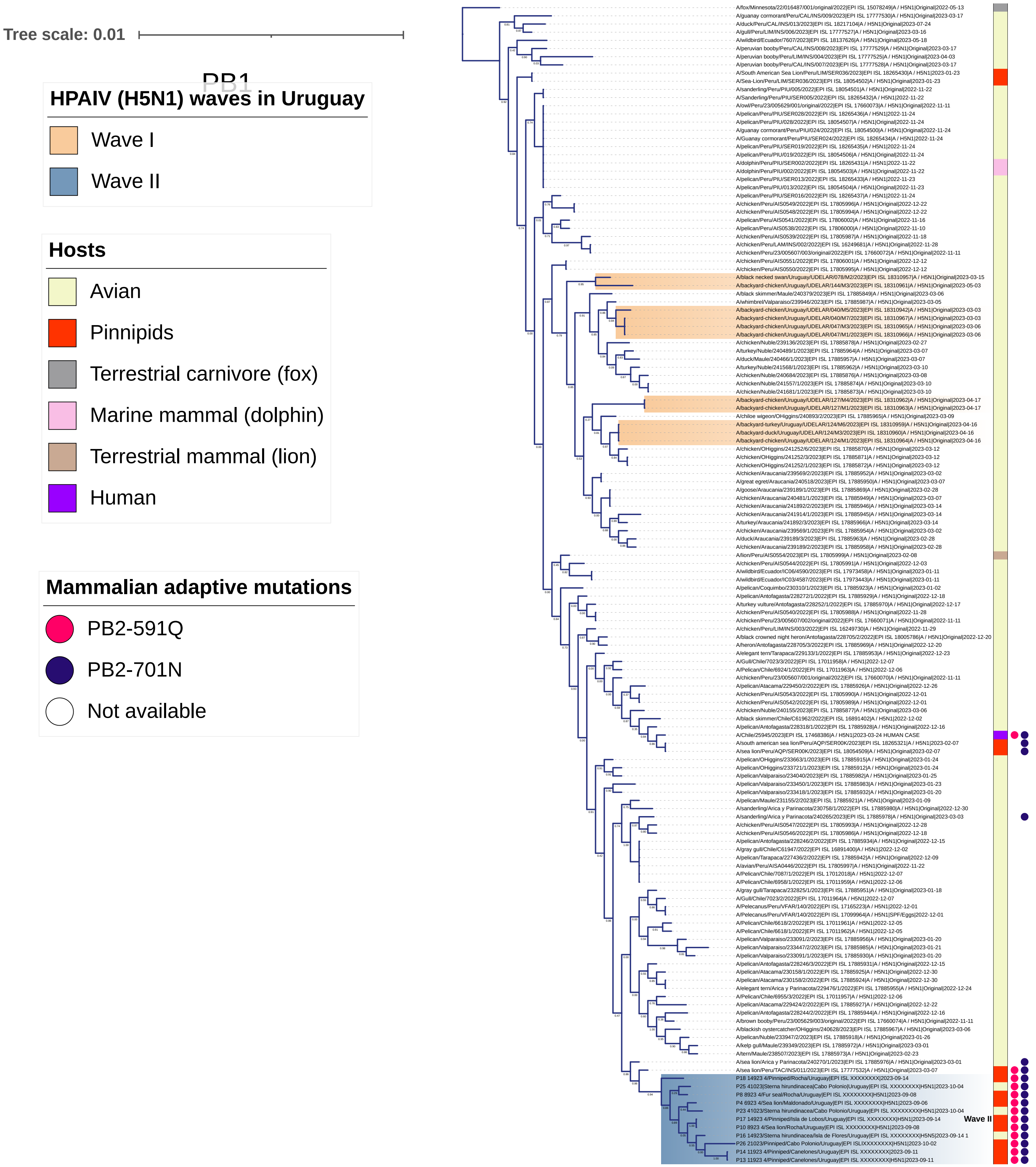

### Appendix Figure 1C (segment 3 PA gene)

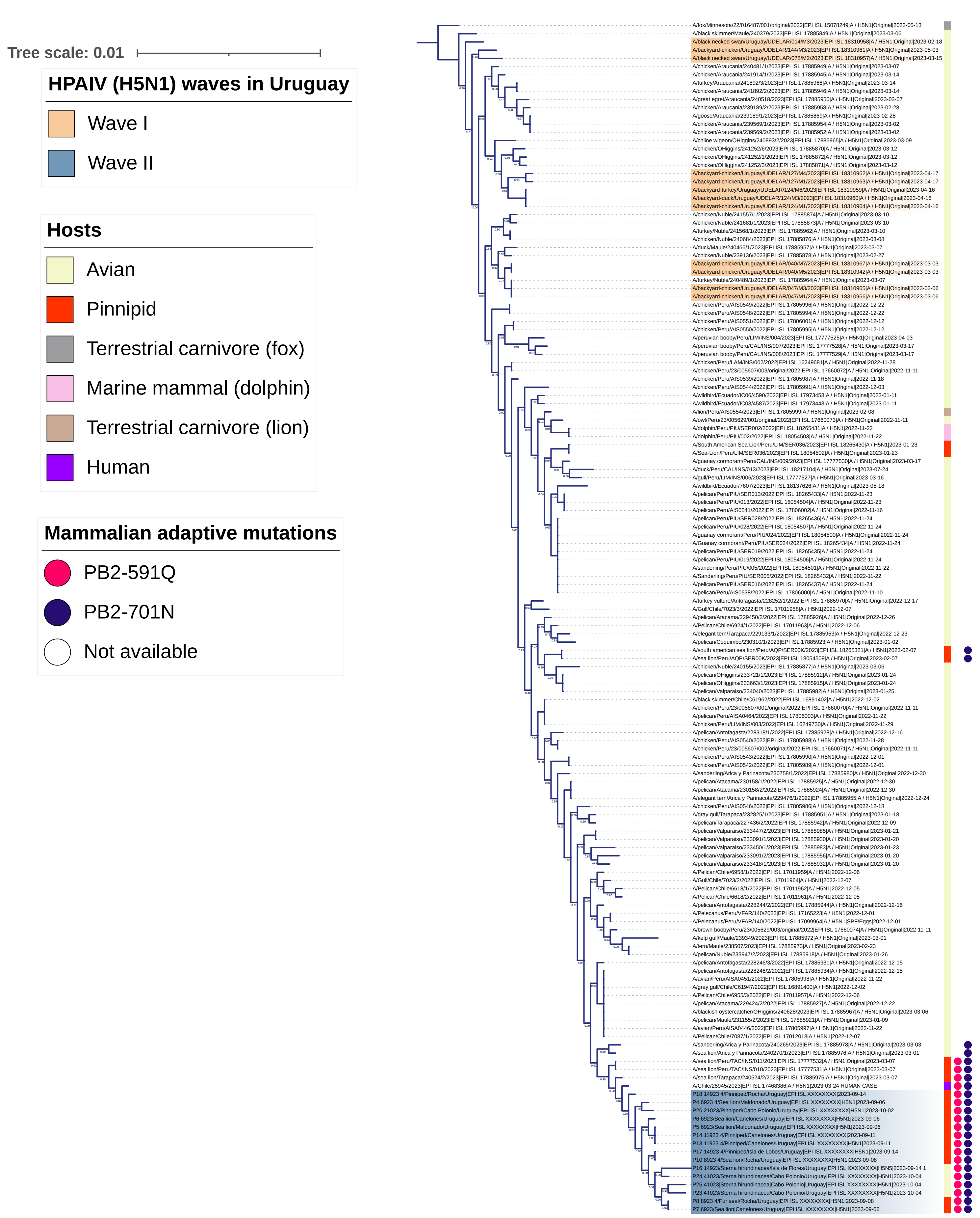

Appendix Figure 1D (segment 4 HA gene)

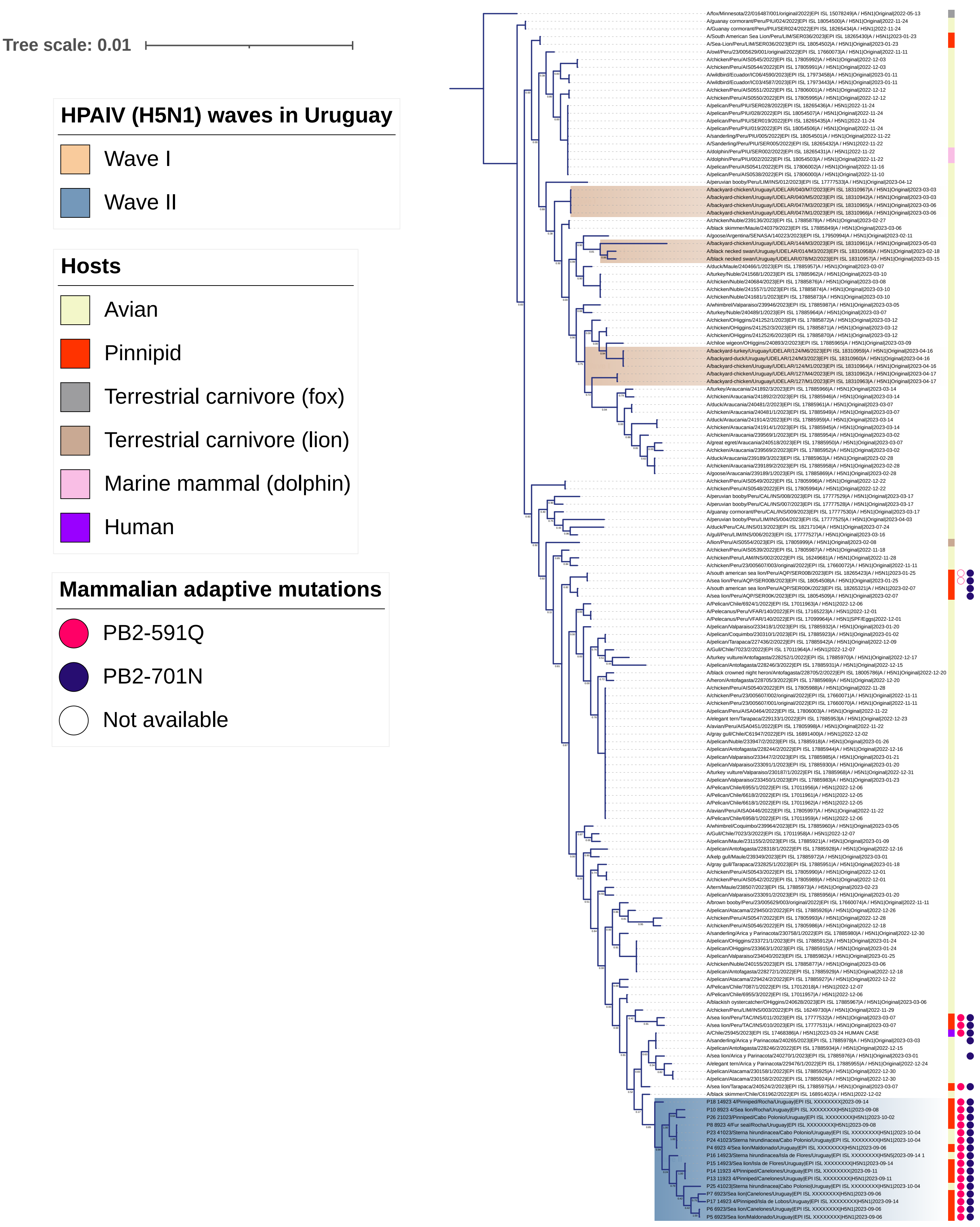

Appendix Figure 1E (segment 5 NP gene)

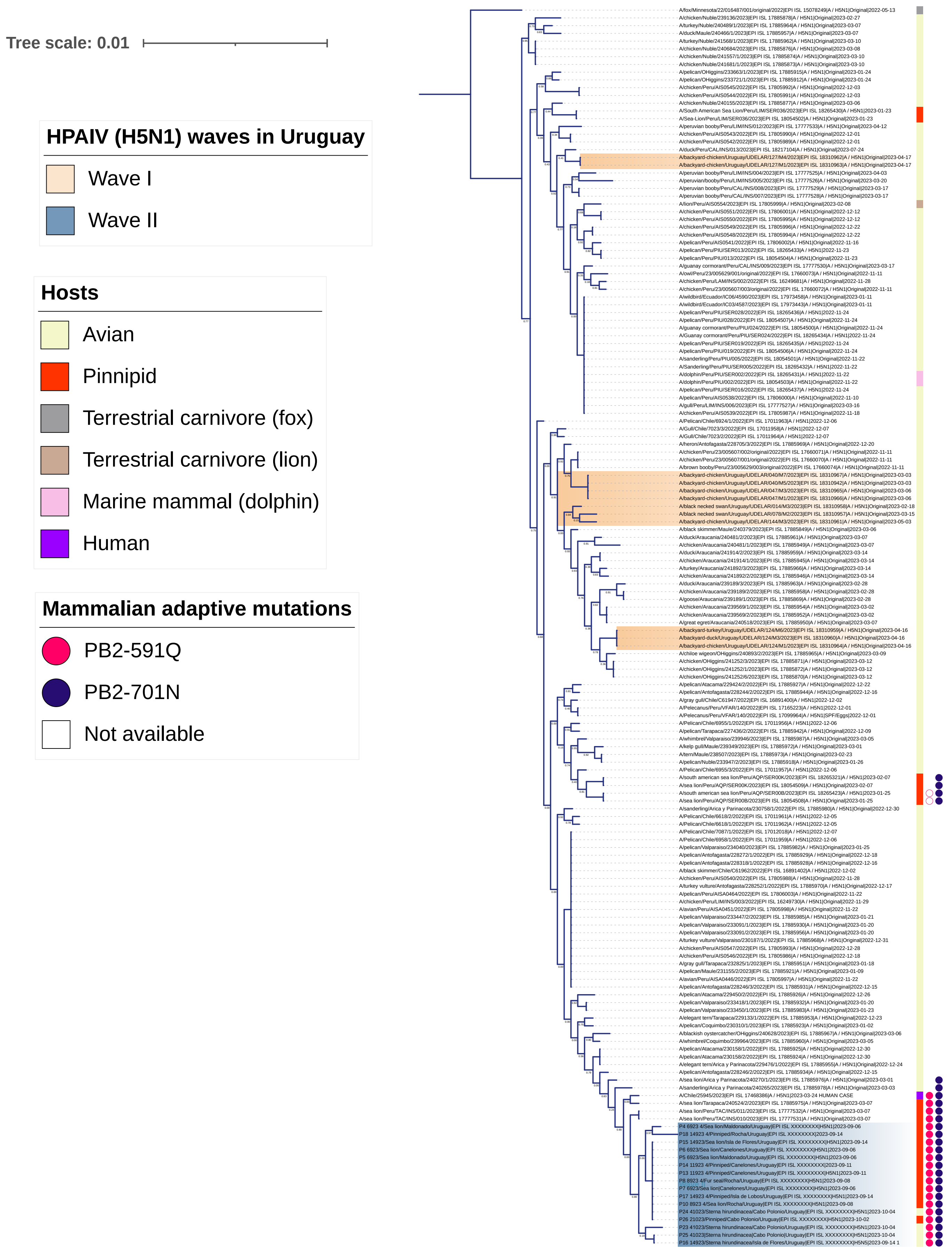

### Appendix Figure 1F (segment 6 NA gene)

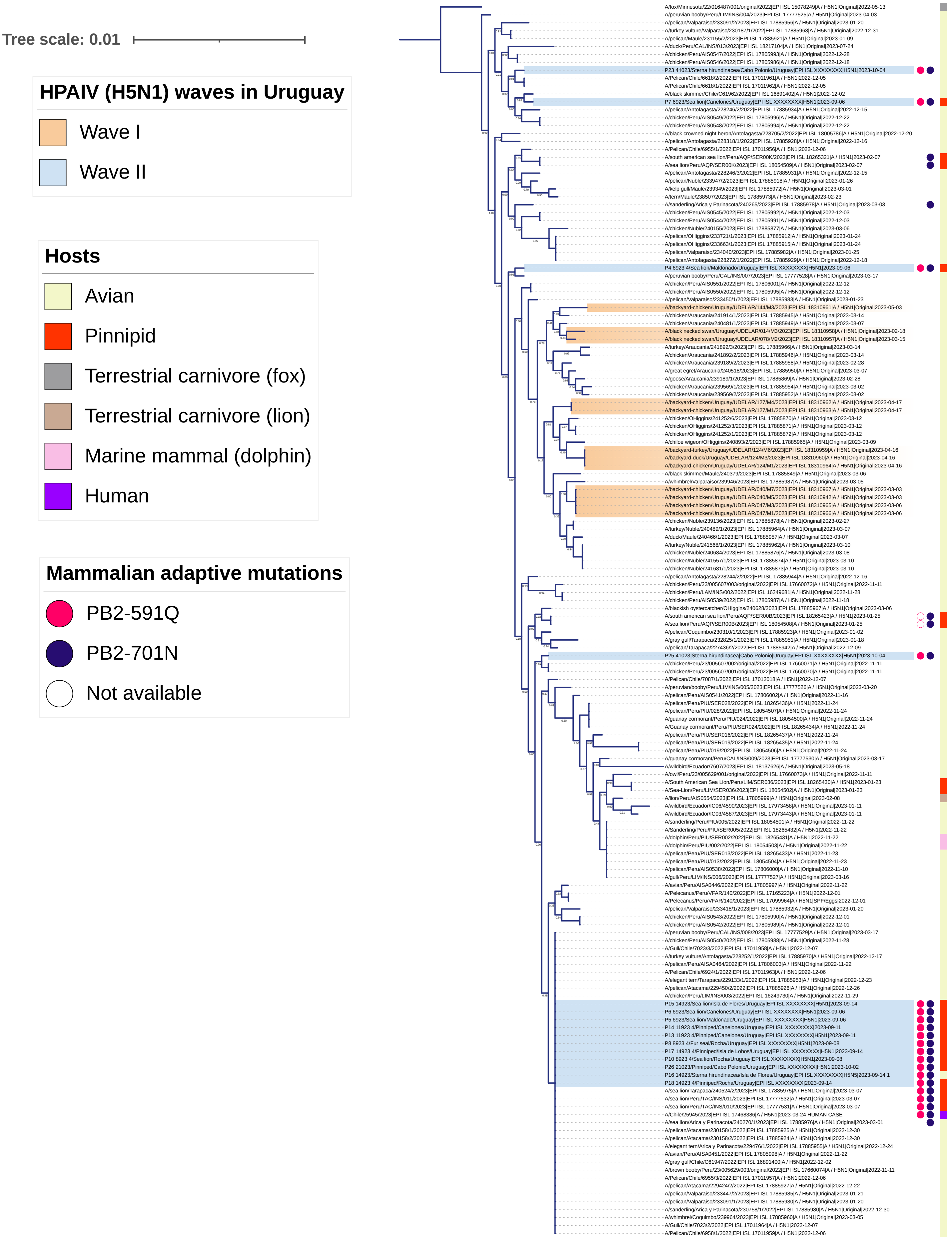

Appendix Figure 1G (segment 7 M gene)

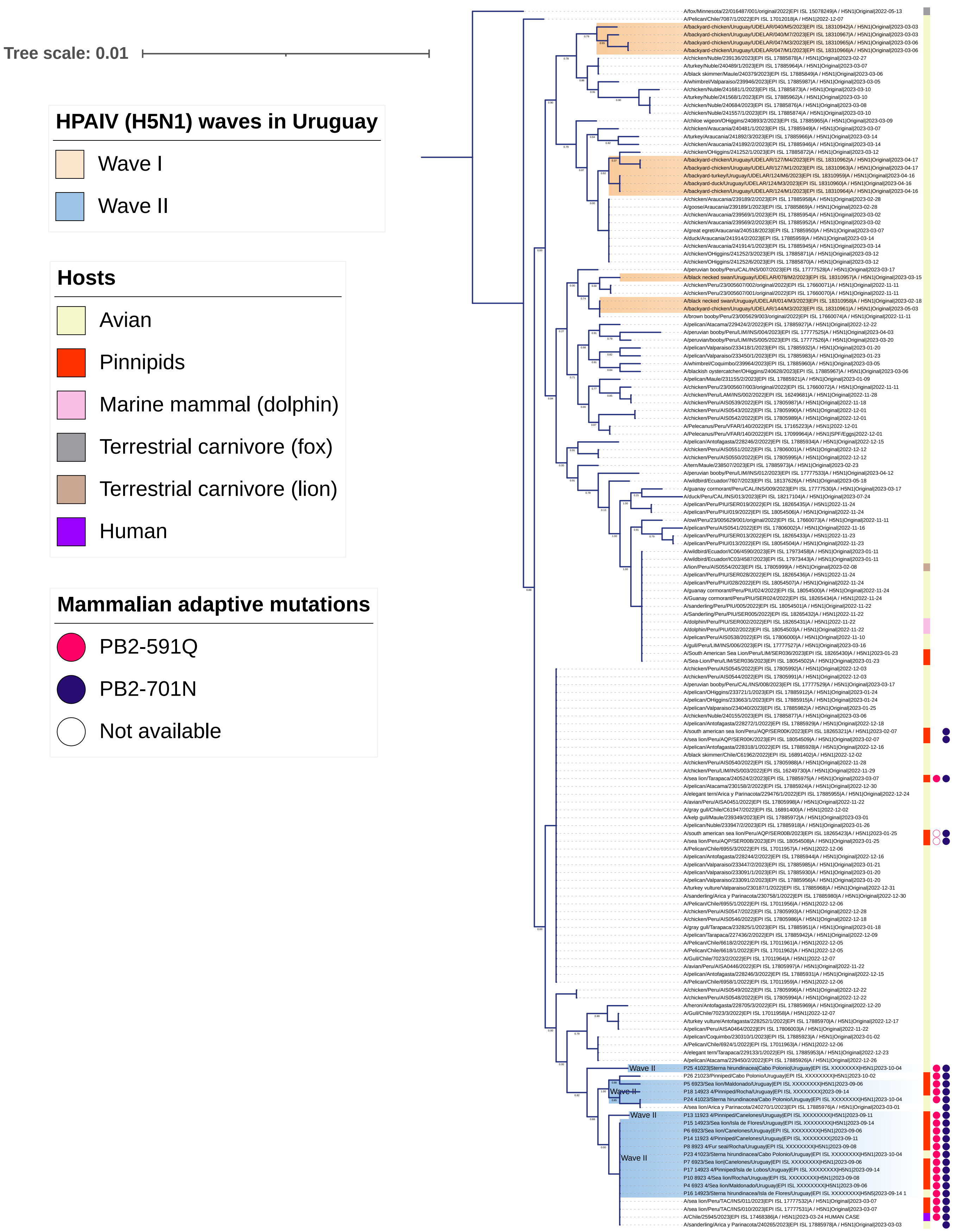

Appendix Figure 1H (segment 8 NS gene)

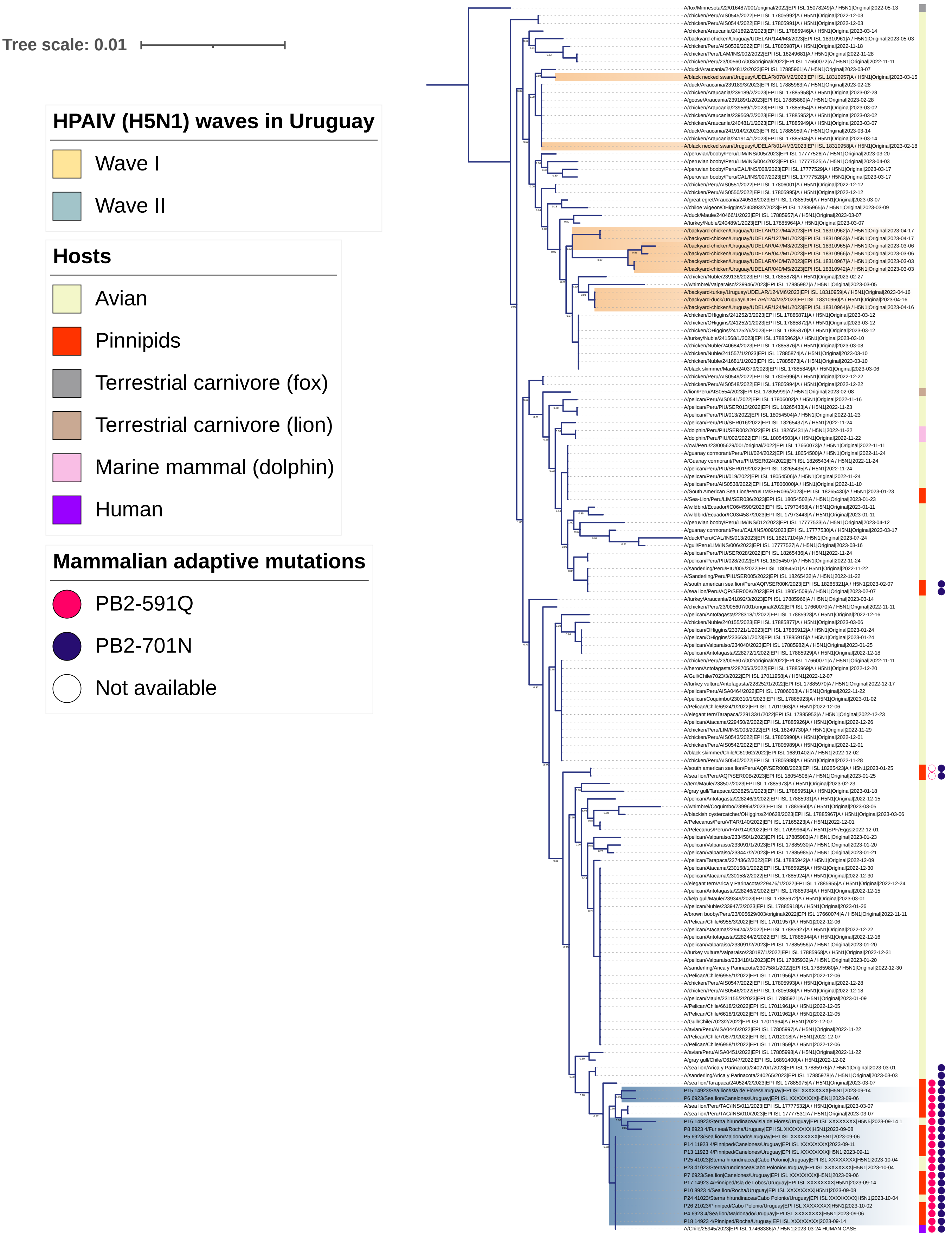

**Appendix Figure 2**: Variation in the distribution of the highly pathogenic avian influenza H5N1 avian influenza virus in South America (Microreact instance including all confirmed cases cataloged by FAO in South America after January 1$^{st}$, 2023). A) Cases until 31 July 2023. Registers showed an initial coastal and an internal spreading involving wild birds, backyard poultry, and outbreaks in commercial farms. B) Cases from 1$^{st}$ August to October 2023. Registers appeared south of the Atlantic coast and spread northward, with fewer cases involving birds and case detections in the interior of South America.

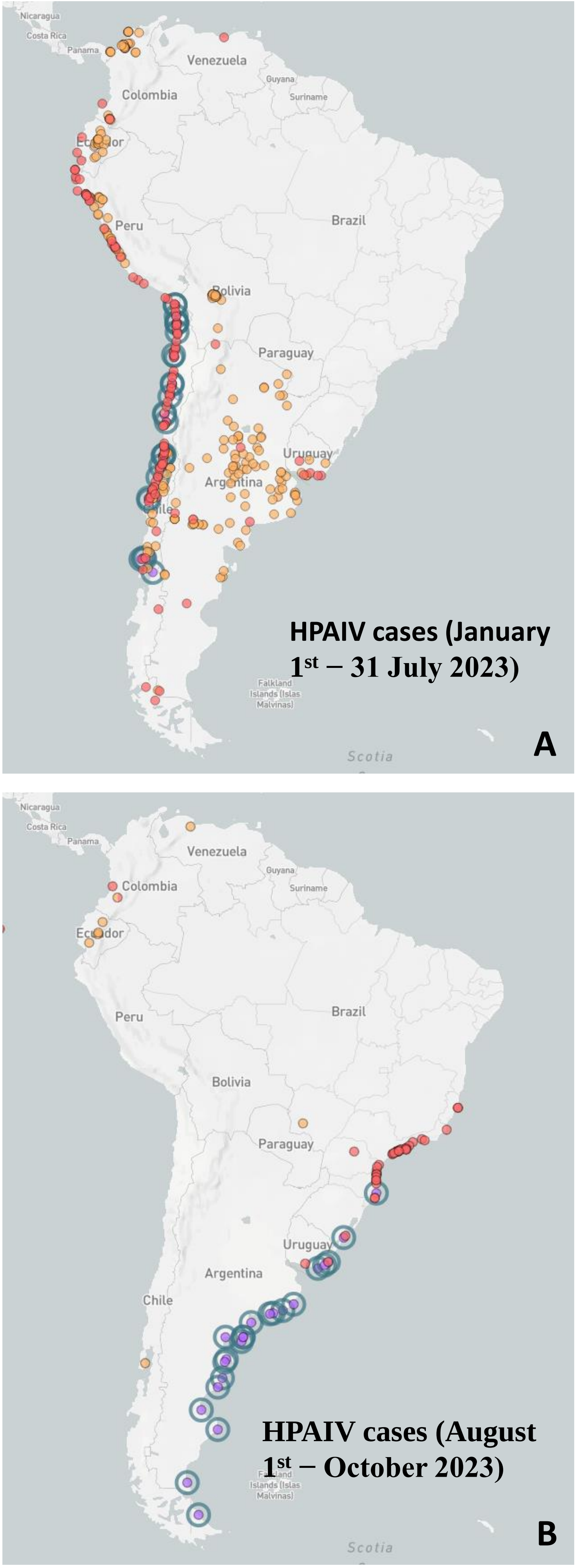


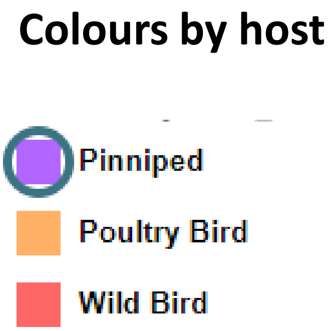


Appendix Figure 2
